# Transcriptomic profiling of *Streptococcus pyogenes* M1T1 strain in a mouse model of necrotizing fasciitis

**DOI:** 10.1101/610139

**Authors:** Yujiro Hirose, Masaya Yamaguchi, Daisuke Okuzaki, Daisuke Motooka, Hiroshi Hamamoto, Tomoki Hanada, Tomoko Sumitomo, Masanobu Nakata, Shigetada Kawabata

**Author notes:** Corresponding author (SK).

## Abstract

*Streptococcus pyogenes* is a major cause of necrotizing fasciitis, a life-threatening subcutaneous soft-tissue infection. At the host infection site, the local environment and interaction between host and bacteria affect bacterial gene-expression profiles, but the *S. pyogenes* gene-expression pattern in necrotizing fasciitis remains unknown. In this study, we used a mouse model of necrotizing fasciitis and performed RNA-sequencing (RNA-seq) analysis of *S. pyogenes* M1T1 strain 5448 by using infected hindlimbs obtained at 24, 48, and 96 h post-infection. The RNA-seq analysis identified 483 bacterial genes whose expression was consistently altered in the infected hindlimbs as compared to their expression under *in vitro* conditions. The consistently enriched genes during infection included 306 genes encoding molecules involved in virulence, carbohydrate utilization, amino acid metabolism, trace-metal transport and vacuolar ATPase transport system. Surprisingly, drastic upregulation of 3 genes, encoding streptolysin S precursor (*sagA*), cysteine protease (*speB*), and secreted DNase (*spd*), was noted in the mouse model of necrotizing fasciitis (log_2_ fold-change values: >6.0, >9.4, and >7.1, respectively). Conversely, the consistently downregulated genes included 177 genes, containing genes associated with oxidative-stress response and cell division. These results suggest that *S. pyogenes* in necrotizing fasciitis changes its metabolism, decreases cell proliferation, and upregulates the expression of major toxins. Our findings could provide critical information for developing novel treatment strategies and vaccines for necrotizing fasciitis.

**Author summary:** Necrotizing fasciitis, a life-threatening subcutaneous soft-tissue infection, principally caused by a *Streptococcus pyogenes*. At infection sites in hosts, bacterial pathogens are exposed to drastically changing environmental conditions and alter global gene expression patterns for survival and pathogenesis. However, there is no previous report about transcriptomic profiling of *S. pyogenes* in the necrotizing fasciitis. Here, we conducted comprehensive gene-expression analyses of *S. pyogenes* in the mouse model of necrotizing fasciitis at three distinct time points during infection. Our results indicated that *S. pyogenes* drastically upregulates the expression of virulence-associated genes and shifts metabolic-pathway usage during infection. The high-level expressions in particular of toxins, such as cytolysins, proteases, and nucleases, were observed at infection sites. In addition, the consistently enriched genes identified here included genes for metabolism of arginine and histidine, and carbohydrate uptake and utilization. Conversely, the genes associated with oxidative-stress response and cell division were consistently downregulated in the mouse model of necrotizing fasciitis. These data will provide useful information necessary for establishing novel treatment strategies (166 words).

## Introduction

*Streptococcus pyogenes* causes diverse human diseases, ranging from mild throat and skin infections to life-threatening invasive diseases such as sepsis, necrotizing fasciitis, and streptococcal toxic-shock syndrome. Streptococcal necrotizing fasciitis cases are clinically characterized by fulminant tissue destruction and rapid disease progression [1]. In a majority of the cases, surgical treatment is required, including amputations, in addition to intensive care. Although this infection has been attracting increasing research and clinical interest, the mortality rate remains high [2, 3]. Investigation of the molecular pathogenesis of *S. pyogenes* in necrotizing fasciitis is expected to lead to the development of novel therapeutic strategies or effective treatments.

*S. pyogenes* typing has been historically conducted on the basis of antigenicity of M protein and T antigen (pilus major subunit). Currently, the sequence typing of the region encoding hypervariable region of M protein has been widely applied to classify this organism into at least 240 *emm* sequence types [4–6] and ~20 T serotypes [7, 8] have been identified. In industrialized societies, *S. pyogenes* serotype M1 (*emm* 1) isolates are considerably more common than other serotypes among invasive cases [6, 9–11], and the M1T1 clone in particular is the most frequently isolated serotype from severe invasive human infections worldwide [12, 13].

At infection sites in hosts, bacterial pathogens are exposed to drastically changing environmental conditions, which include the host cells, tissues, and immune response, as compared to laboratory growth conditions. However, no previous report has compared *in vivo* and *in vitro* transcriptome of *S. pyogenes*

Several comprehensive *in vitro* analyses of *S. pyogenes* gene expression performed using microarray or RNA-sequencing (RNA-seq) approaches and revealed the roles of *S. pyogenes* virulence-related regulators [14], such as CovRS [15, 16] and CcpA [17–19]. The transcriptome profile of *S. pyogenes* from a mouse soft-tissue infection, which was obtained using microarray analysis, indicated that *S. pyogenes* MGAS5005 (serotype M1) upregulated genes that are involved in oxidative-stress protection and stress adaptation [20]. The results of another microarray analysis on *S. pyogenes* MGAS5005 (serotype M1) demonstrated downregulation of glycolysis genes and induction of genes involved in amino acid catabolism and of several types of virulence genes in human blood [21]. These reports suggest that *S. pyogenes* changes its expression levels of virulence factor and metabolic pathways to adapt to the host environment.

Comprehensive understanding of bacterial transcriptomes *in vivo* will facilitate research aimed at developing therapeutic strategies or effective vaccine antigens. Although transposon-directed insertion-site sequencing using cynomolgus macaque model of necrotizing fasciitis was conducted recently [22], this method cannot assess the expression level of genes. In addition, transcriptome analysis of *S. pyogenes* in necrotizing fasciitis have not been performed to date. Here, we investigated the transcriptome profiling of *S. pyogenes* M1T1 strain 5448 by using a mouse model of necrotizing fasciitis, from the acute phase to the elimination phase, and we identified genes whose expression is consistently altered throughout the infection period. This new information may shed light on the development of novel therapeutic strategies for the infection.

## Results

### Establishing a technique for high-yield purification of bacterial RNA from mouse tissue

We used a mouse necrotizing fasciitis model as described previously, with minor modifications [23]. At 24 and 48 h after infection, the mouse model of necrotizing fasciitis was histologically similar to human necrotizing fasciitis in terms of tissue necrosis, the infection spread along the fascial planes, inflammatory-cell infiltration, hemorrhage and ulceration [24, 25] (Fig 1A and 1B). Extensive scab formation was detected at 48 h post-infection and elimination of pus from infected hindlimbs was observed at 96 h post-infection. At 96 h after infection, the weight of the mice tended to recover, and the bacterial burden at the infected site also decreased (S1 Fig). Therefore, we collected infected hindlimb samples from the 24, 48, and 96 h groups. To obtain bacterial RNA from infected tissues, we established a suitable protocol by using two types of beads (Fig 1C); this method allows us to remove most mouse RNA from samples and obtain high-yield purification of bacterial RNA (Fig 1D).

**Fig 1.**
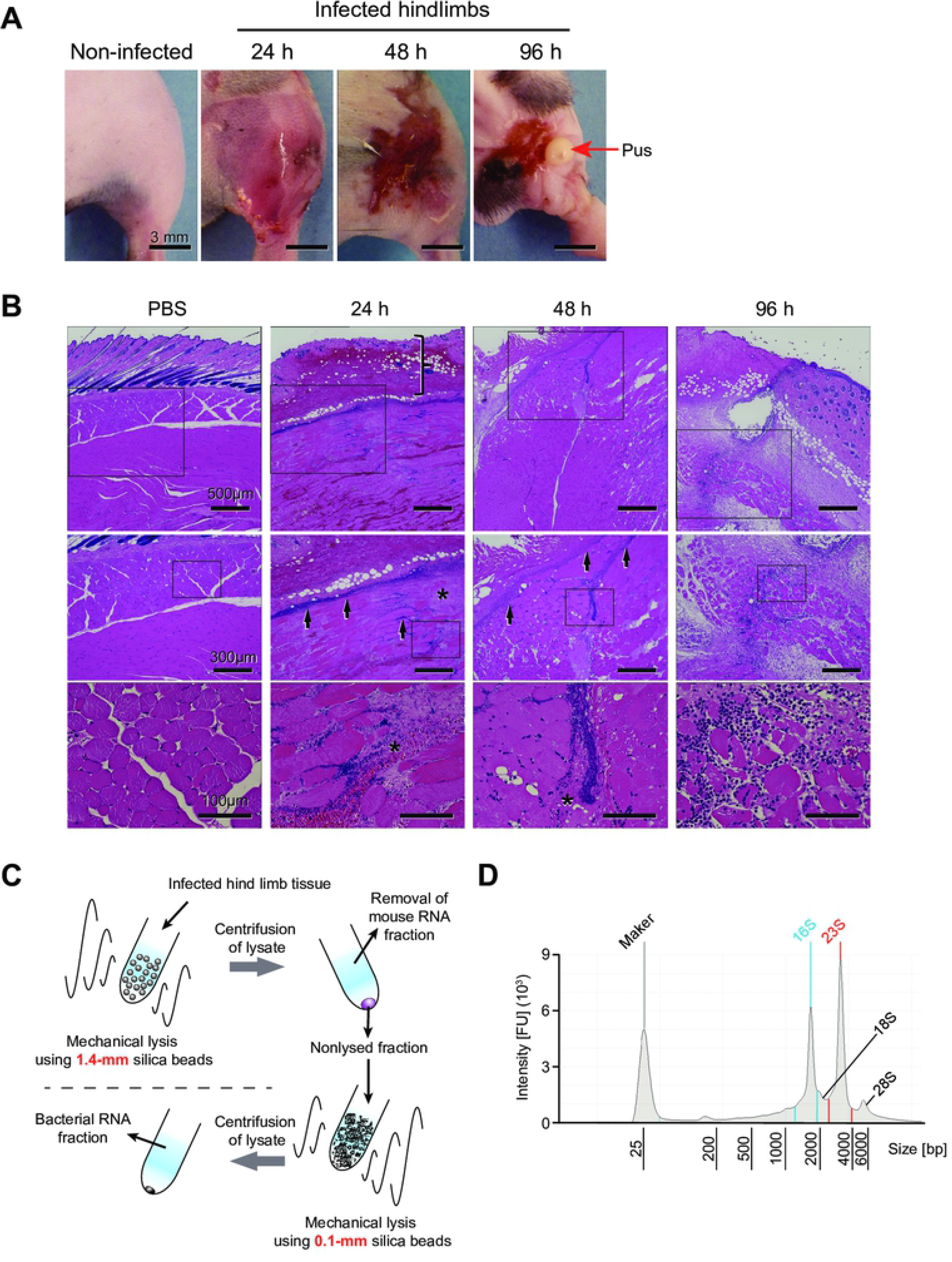
Features of a mouse model of necrotizing fasciitis and workflow of bacterial RNA isolation. (A) Representative images of infected hindlimbs after inoculation with *S. pyogenes* M1T1 strain 5448. (B) Histopathological features of the mouse model of necrotizing fasciitis. Hematoxylin and eosin staining of infected lesions at the indicated time points is shown, and higher-magnification images of the selected areas of the same sections are presented. At 24 h after infection, the skin shows erosion of the epidermis and edematous thickening of the dermis (vertical bracket), as well as sparse inflammatory-cell infiltration. At 24 and 48 h after infection, marked necrosis (asterisks) is observed, as is the presence of bacteria concentrated primarily along the major fascial planes (arrows) in the infected deep soft tissue. At 96 h after infection, sufficient inflammatory-cell infiltration and elimination of pus (red arrows) from infected hindlimbs are observed. (C) Workflow of bacterial RNA isolation. At first, tissues were lysed with 1.4-mm silica spheres and mouse RNA fraction was removed after centrifugation. Next, pellets were lysed with 0.1-mm silica spheres and centrifuged to obtain bacterial RNA fraction. (D) Representative bioanalyzer profile of total RNA isolated from an infected hindlimb; 16S and 23S: bacterial rRNA peaks; 18S and 28S: mouse rRNA peaks.

### Similar gene-expression patterns of *S. pyogenes* at three distinct time points in infected hindlimbs

We performed RNA-seq analysis on *S. pyogenes* isolated from infected hindlimbs at 24, 48, and 96 h post-infection. RNA-seq data from *S. pyogenes* during the exponential growth phase in THY medium (Todd-Hewitt broth plus yeast extract) were defined as the control. To assess the global gene-expression profiles of the samples, we performed principal component analysis (PCA) (Fig 2A), hierarchical clustering analysis (S2 Fig), and k-means clustering (Fig 2B) by using the RNA-seq data. In PCA and hierarchical clustering analysis, bacterial RNA expression patterns of THY culture samples formed a cluster and the samples from the infected tissues were well separated. The 24 h_1 sample showed a global gene-expression profile that was distant from the profiles of other samples, whereas the heatmap of k-means clustering showed that the gene-expression profile of 24 h_1 was at least partially similar to that of samples from the infected tissues as indicated in Clusters A and B. The k-means clustering further suggested that most samples from infected hindlimbs show changes in global mRNA transcript patterns in opposite directions as compared to the THY group.

**Fig 2.**
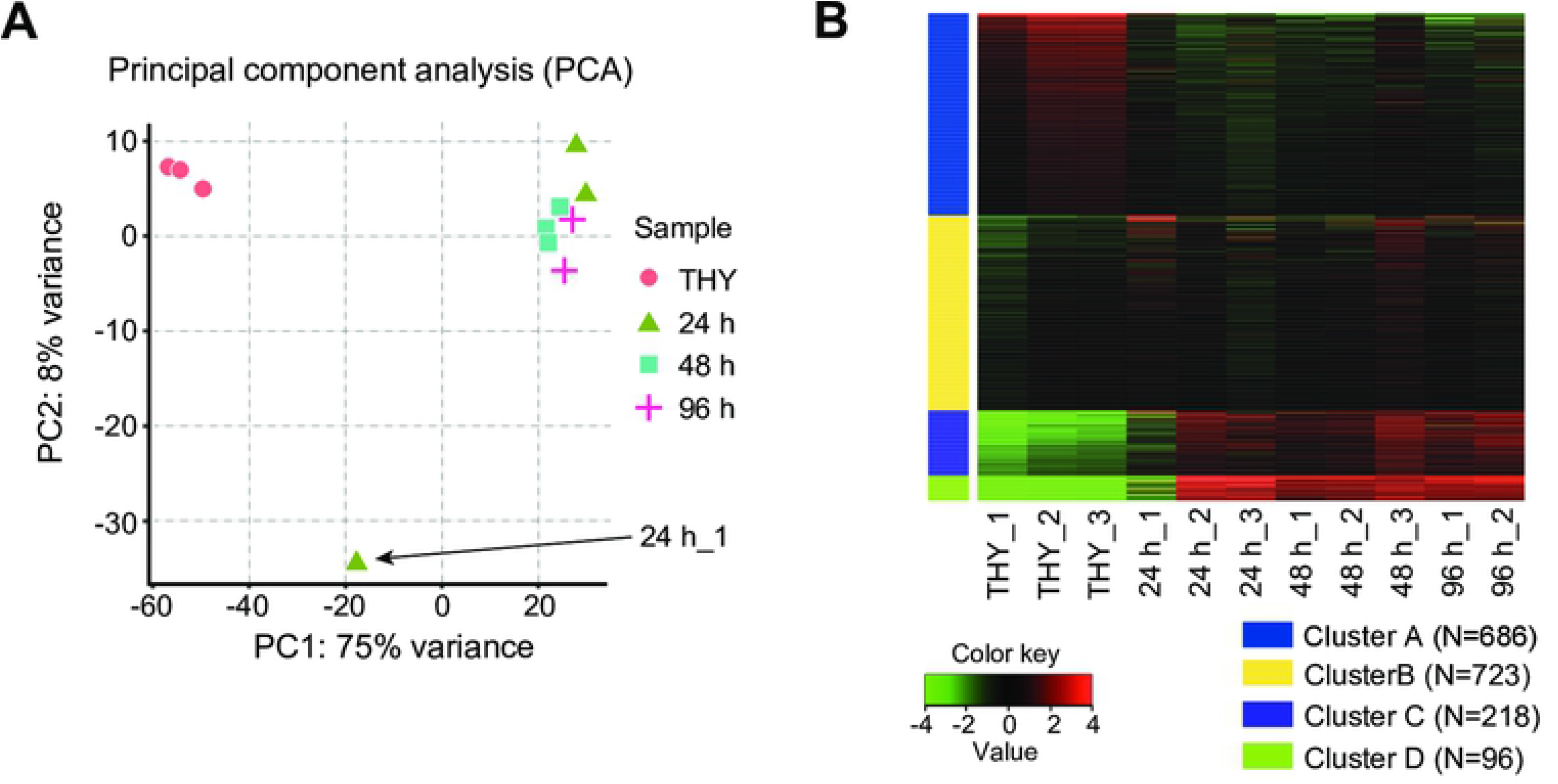
RNA-seq global reports. (A) Principal component analysis (PCA) plot of RPKM data from RNA-seq dataset. (B) Heatmap of k-means clustering of all genes (1,723 genes) in all samples (k = 4). The number of expressed genes in each cluster is indicated. The color key indicates Z-scores, which display the relative values of all tiles within all samples: green, lowest expression; black, intermediate expression; red, highest expression. Bacterial RNA-seq data at 24, 48, and 96 h post-infection were defined as 24-h group (24 h_1, 24 h_2, 24 h_3), 48-h group (48 h_1, 48 h_2, 48 h_3), and 96-h group (96 h_1, 96 h_2), respectively. Bacterial RNA-seq data of THY culture samples were defined as the control and named THY group (THY_1, THY_2, THY_3).

### Consistently altered 483 bacterial genes at three time points in mouse necrotizing fasciitis model

Differentially expressed genes (DEGs; absolute log_2_ fold-change > 1 and adjusted P < 0.1) were detected between *S. pyogenes* in infected tissues and in THY broth (Fig 3A). In S1 Dataset, we provide the DEG details and list the following information for all the genes: gene ID, gene name, the gene-associated function, log_2_ fold-change, adjusted P value, and reads per kilobase per million mapped reads (RPKM) value. In comparisons of 48 h vs 24 h groups and 96 h vs 24 h groups, no DEGs were detected (Fig 3A), and only 4 DEGs were detected between 96 h and 48 h groups (S1 Dataset). These results indicate that *S. pyogenes* expresses similar genes at the three time points in the mouse model of necrotizing fasciitis. To identify the genes that are consistently enriched or downregulated in the mouse necrotizing fasciitis model, we drew Venn diagrams by using the DEGs from the comparisons of 24 h and THY groups, 48 h and THY groups, and 96 h and THY groups (Fig 3B); 28.0% of all 1,723 genes, 483 genes, were identified as consistently altered bacterial genes in infected hindlimbs (S2 Dataset). Among the 483 genes, 306 and 177 genes were upregulated and downregulated, respectively, at all three time points.

**Fig 3.**
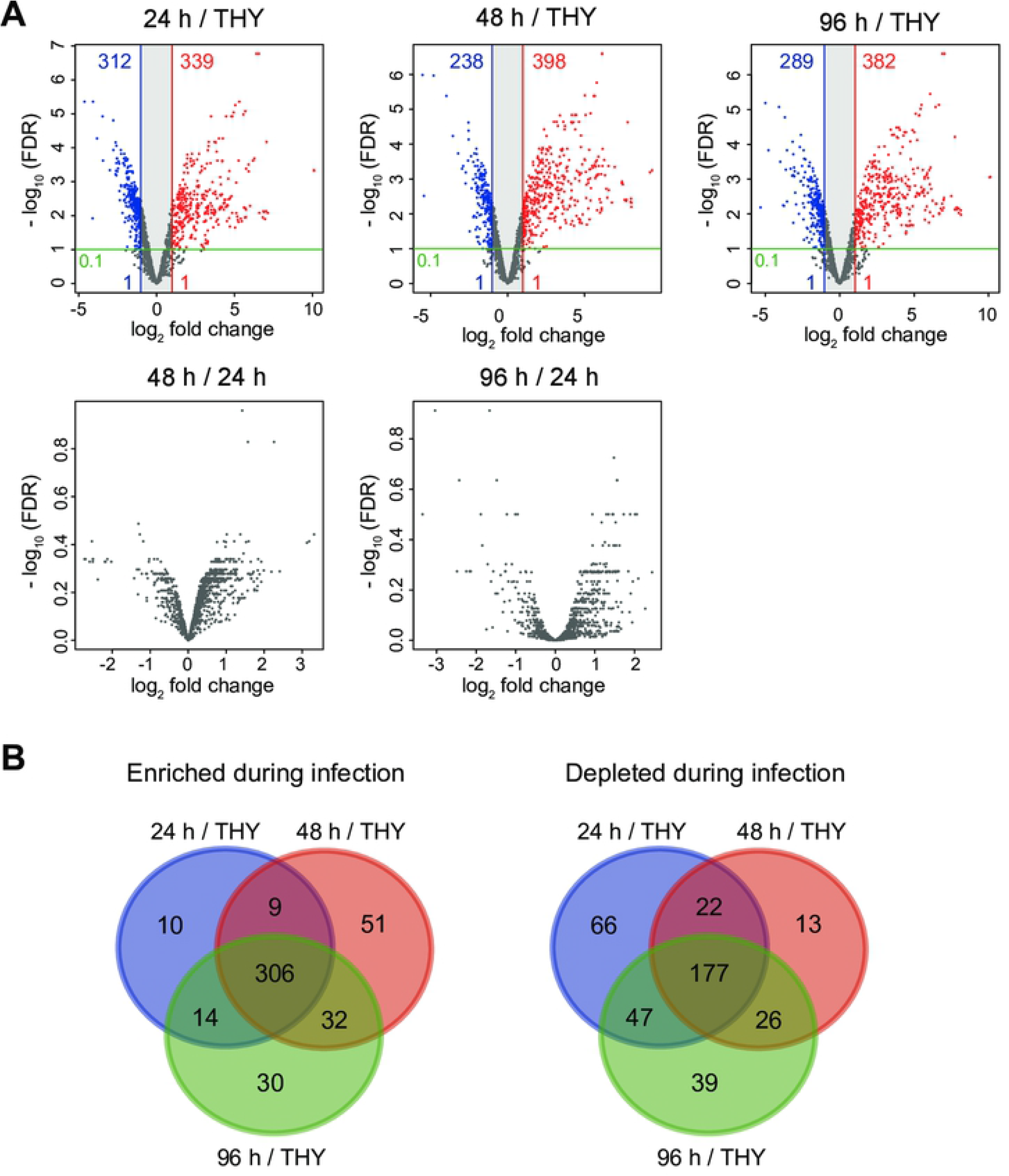
Differentially expressed genes showing consistent alteration in hindlimbs at each infection phase. (A) Volcano plots showing gene-expression differences under the comparison conditions indicated in each figure. Colored circles: significantly upregulated (red) and downregulated (blue) genes (absolute log_2_ fold-change > 1; adjusted P < 0.1). (B) Three-way Venn diagram illustrating bacterial genes that are consistently altered during infection (24 h vs THY, 48 h vs THY, 96 h vs THY): relative to THY condition, 306 transcripts were consistently enriched *in vivo* (log_2_ fold-change > 1; adjusted P < 0.1) and 177 transcripts were consistently downregulated *in vivo* (log_2_ fold-change < −1; adjusted P < 0.1).

### Marked upregulation of genes encoding virulence factors

The consistently enriched genes featured a high proportion of genes encoding virulence factors, such as genes for cytolysins (*sagA-I, slo*), nucleases (*spd, spd3, sdaD2*), cysteine protease (*speB*), factors involved in immune evasion (*endoS, spyCEP, scpA*, sic), superantigens (*speA, smeZ*), and adhesins (*fbaA*, *lbp, emm*) (Table 1; S2 Dataset). Surprisingly, the RPKM values of genes encoding streptolysin S precursor (*sagA), speB*, SpeB–inhibitor-encoding gene (*spi*), and *spd* were extremely high and were consistently ranked within the top four (S1 Dataset; Fig 4). As compared with their expression under the THY condition, *sagA*, *speB*, *spi*, and *spd* were expressed in mouse necrotizing fasciitis at the following levels (respectively): log_2_ fold-change = >6.0, >9.3, >9.4, and >7.1. By contrast, the gene encoding macroglobulin-binding protein (*grab*) was markedly downregulated at all three time points. Lastly, hyaluronic acid synthesis operon (*hasABC*) was significantly upregulated in the 48 h and 96 h groups, but no significant difference was detected in the 24 h group.

**Fig 4.**
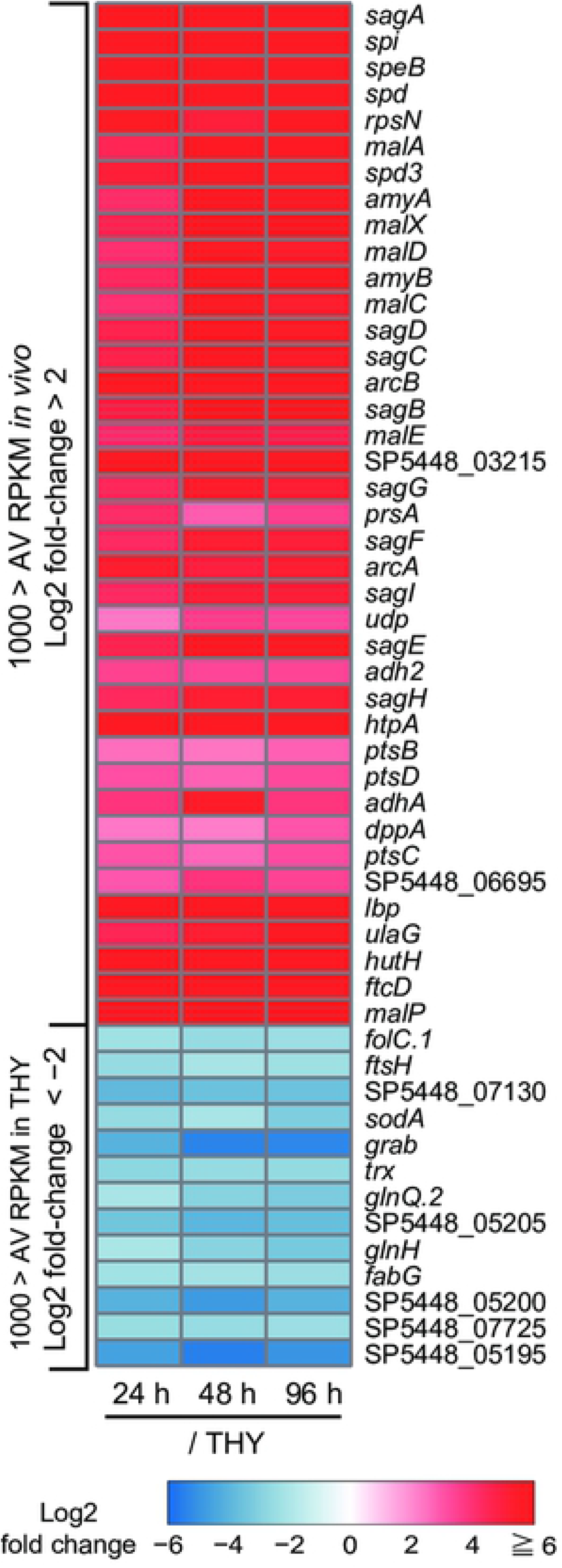
Heatmap of significantly altered genes *in vivo*. Heatmap of consistently and significantly enriched genes (log_2_ fold-change > 2, average RPKM *in vivo* > 1,000) or downregulated genes (log_2_ fold-change < −2, average RPKM in THY > 1,000); color scale indicates enrichment (red) and depletion (blue) during infection. Values represent the log_2_ fold-change between indicated conditions, and genes are arranged in descending order of expression level (average of RPKM values *in vivo*). AV RPKM, average RPKM.

**Table 1.**
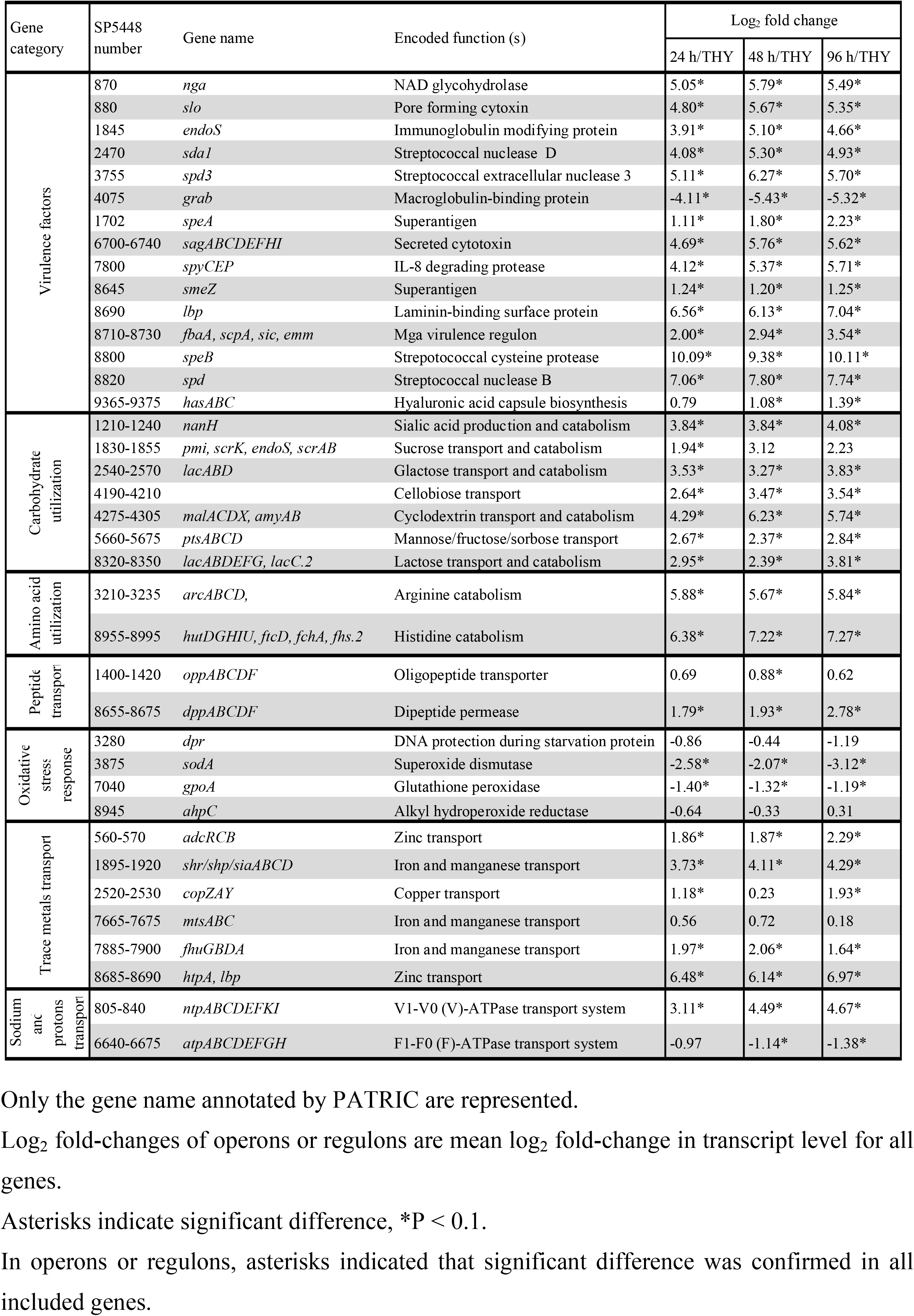
The expression levels of selected genes/operons/regulons.

### Upregulation of carbohydrate uptake and utilization genes

The consistently enriched genes also included most genes encoding ATP-binding cassette (ABC) transporters or phosphoenolpyruvate-phosphotransferase system (PTS) molecules responsible for carbohydrates transport (Fig 5; Table 1; S2 Dataset). In the glycolysis pathway, the expression of *pgk* (encoding phosphoglycerate kinase) and *eno* (encoding enolase) showed a slight decrease, but the RPKM values of these genes consistently remained at >1,500. Despite the sufficient expression of glycolysis-system molecules in the infected hindlimbs, the carbohydrate transport systems exhibited an overall increase. Shelburne *et al*. reported that the carbon catabolite protein CcpA upregulates the expression of most operons encoding transporters of carbohydrates, such as glucose, lactose, maltodextrin, mannose, fructose, cellobiose, lactose, galactose, and sialic acid, under glucose-limiting conditions [18]. Moreover, our results indicated that the genes encoding phosphocarrier protein (*ptsH*) and its kinase (*ptsK*) were consistently downregulated (Fig 5). When Gram-positive bacteria are in the presence of glucose, phosphocarrier protein (HPr) is phosphorylated at Ser46 by its kinase, HprK, which allows phosphorylated HPr to dimerize with CcpA; the dimerized proteins then bind to catabolite-response elements present in promoter sequences and elicit carbon catabolite repression [26]. These findings raise the possibility that *S. pyogenes* is relieved from carbon catabolite repression in the mouse model of necrotizing fasciitis.

**Fig 5.**
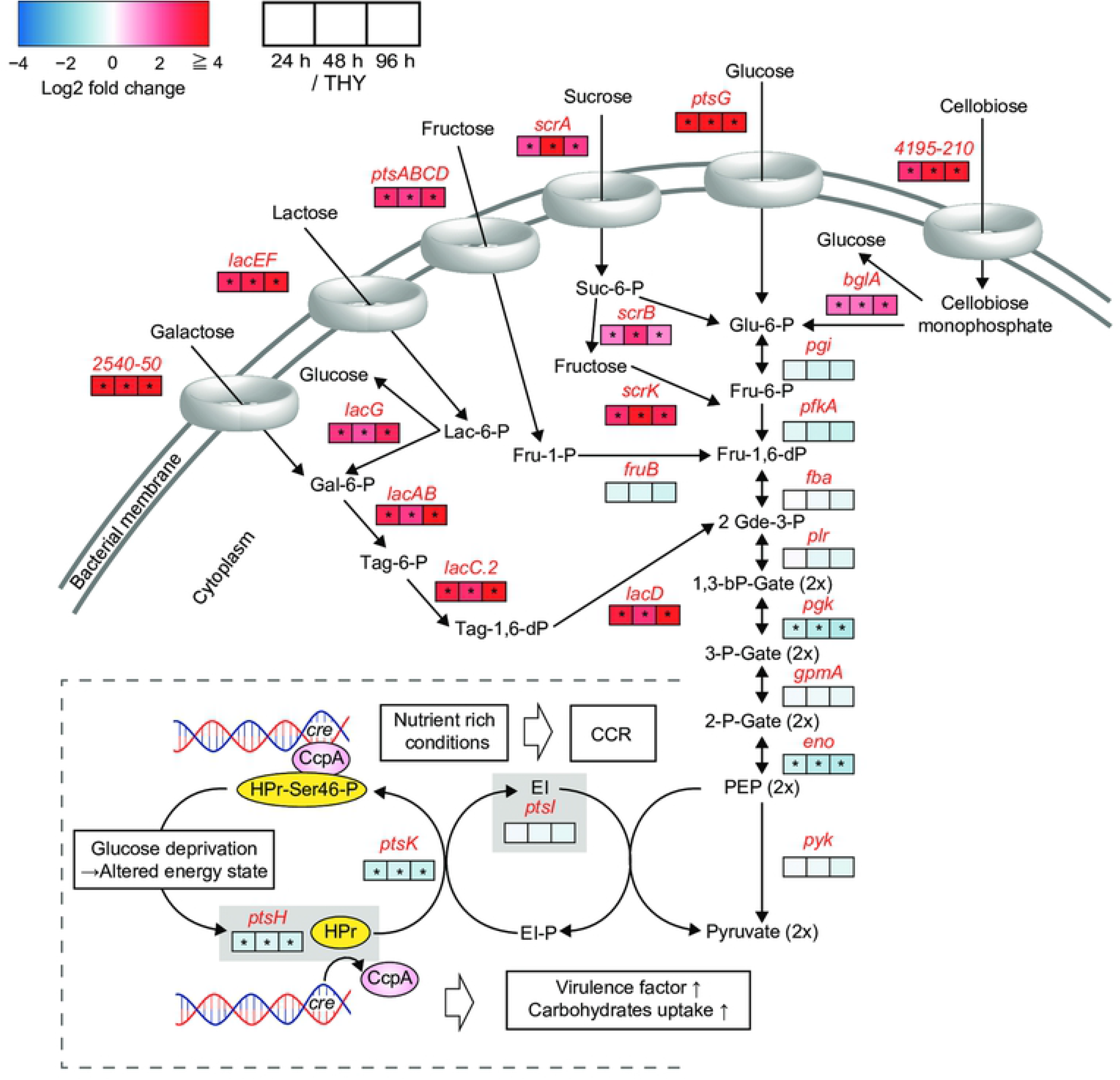
Central carbon metabolism and catabolite control protein CcpA. The pathway shown was constructed based on BioCyc database for *S. pyogenes* MGAS5005. Metabolite names are written in black and gene names are written in red. The log_2_ fold-changes of the 24-h group (left), 48-h group (center), and 96-h group (right) with respect to THY group are indicated in the color-scaled boxes. Color scale indicates enrichment (red) and depletion (blue) during infection. Asterisks indicate significant difference: *P < 0.1. In operons or regulons, asterisks indicate that significant difference was confirmed in all included genes. Log_2_ fold-change values of operons or regulons are mean log_2_ fold-changes in transcript levels for all genes. The phosphocarrier protein HPr (*ptsH*) is phosphorylated at Ser46 by the kinase HPrK (*ptsK*) through the cytoplasmic enzyme EI (*ptsI*), which allows HPr-Ser46-P to dimerize with the carbon catabolite protein CcpA and elicit carbon catabolite repression by binding to catabolite-response elements in promoter sequences [26].

### Drastically enhanced arginine and histidine metabolism in infected hindlimbs

*S. pyogenes* is auxotrophic for at least 15 amino acids [27]. The consistently enriched genes identified here included operons for metabolism of arginine (*arcABCD*), histidine (*hutDGHIU, ftcD, fchA, fhs.2*), and serine (*salB*) (Table 1; Fig 4; Fig 6; S2 Dataset). Conversely, the bacterial genes encoding proteins for isoleucine metabolism (*bcaT, acoC*) were consistently downregulated in the infected hindlimbs (Fig 6). The operon for the dipeptide transporter *dppABCDF*, which is involved in the uptake of essential amino acids [28], was also upregulated in the infected hindlimbs (Table 1), and the expression of *dppA*, which encodes a dipeptide-binding protein, was remarkably enhanced (log_2_ fold-change > 2.25) (Fig 4; S2 Dataset).

**Fig 6.**
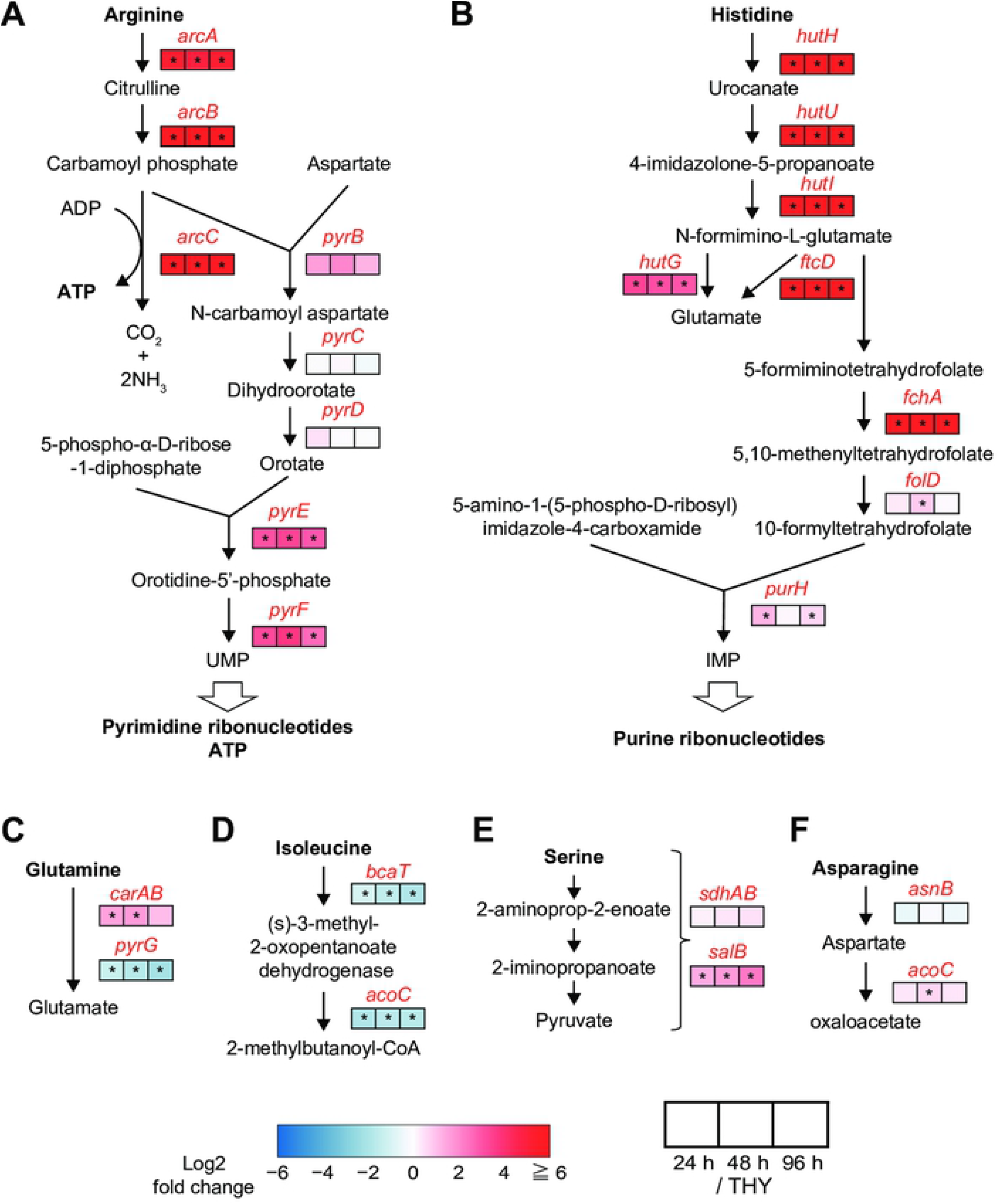
Significant enhancement of arginine and histidine metabolic pathways. (A) Arginine deiminase and pyrimidine nucleotide *de novo* synthesis pathways. (B) Histidine degradation and purine nucleotide *de novo* biosynthesis pathways. (C) Glutamine, (D) isoleucine, (E) serine, and (F) asparagine degradation pathways. Pathways were constructed based on BioCyc database for *S. pyogenes* MGAS5005. Metabolite names are written in black and gene names are written in red. The log_2_ fold-changes of 24-h group (left), 48-h group (center), and 96-h group (right) with respect to THY group are indicated in color-scaled boxes. Color scale indicates enrichment (red) and depletion (blue) during infection. Asterisks indicate significant difference: *P < 0.1. UMP, uridine monophosphate; IMP, inosine monophosphate.

The mean fold-changes in the transcript levels (i.e., the mean log_2_ fold-change values) for all genes in the operons for arginine and histidine metabolism were >5.67 and >6.38, respectively. In *S. pyogenes*, the arginine deiminase pathway (*arcABCD*) is reported to supplement energy production, help protect against acid stress, and compete with arginine-dependent NO production by host cells in the subcutaneous layer [29]. Another critical role of arginine metabolism is to serve as the source of uridine monophosphate (Fig 6A), whereas histidine metabolism is connected to the synthesis of inosine monophosphate (Fig 6B). These functions cooperate with pyrimidine and purine metabolism for the synthesis of DNA and RNA. The consistently enriched genes also included genes for pyrimidine and purine metabolism (S2 Dataset). These results suggest the possibility that bacterial synthesis of nucleic acids is active in infected hindlimbs, although we also observed the repression of certain genes related to cell division, such as genes encoding cell-division proteins (*ftsA*, *ftsZ, ftsH*), amino acid ligases (*murD, murG*), phospho-N-acetylmuramoyl-pentapeptide transferase (*mraY*), and ribonuclease III (*rnc*) (Fig 4; S1 and S2 Datasets).

### Host-induced bacterial stress responses

Genes encoding superoxide dismutase (*sodA*) and glutathione peroxidase (*gpoA*) were consistently downregulated in the infected hindlimbs (Table 1; S2 Dataset). SodA and GpoA act to neutralize endogenous and exogenous peroxides, which contributes to detoxification of reactive oxygen species *in vitro* [30, 31]. Our results suggest that *S. pyogenes* is not exposed to substantial oxidative stress in the infected hindlimbs as compared to the stress encountered during the aerobic growth.

Transition metals are involved in several crucial biological processes in pathogens that are necessary for the pathogens to survive, proliferate, and cause diseases in their environmental niche. In *S. pyogenes*, contributions to virulence are made by the homeostasis of metals, including iron, manganese [32], and zinc [33], whereas hosts exploit this phenomenon and combat invading pathogens by restricting the availability of essential metals by using transferrin (iron), lactoferrin (iron), and calprotectin (manganese and zinc) [34]. Here, *S. pyogenes* was found to upregulate genes involved in iron and manganese transport (*shr/shp/siaABCD, fhuGBDA*) and zinc transport (*adcRCB, htpA, lbp*) in the mouse model of necrotizing fasciitis (Table 1; S2 Dataset).

### Altered expression of virulence-related transcriptional-regulator genes

Expression of most virulence genes in *S. pyogenes* is under the control of two-component signal transduction systems (TCSs) and transcriptional activators/repressors [14]. Although phosphorylation is recognized as a key modification by which regulators exert regional transcriptional control [26, 35, 36], the alternation of regulator gene-expression levels could also influence the degree of regulation.

*S. pyogenes* showed altered expression of several genes encoding virulence-related regulators (Fig 7): The consistently enriched genes included TCS *trxSR* operon and genes encoding carbohydrate-sensitive regulators (*lacD.1, ccpA*), a member of the RofA-like protein type family of stand-alone virulence-related regulators (*rivR/ralp4*), and maltose repressor (*malR)*. Conversely, the only consistently downregulated regulator gene was the gene encoding streptococcal regulator of virulence (srv), although certain other regulators also tended to show downregulation, including the genes for CovRS (*covRS*), the metabolic-control regulator VicRK (*vicRK*), metalloregulator (*mtsR/scaR*), and RofA regulator (*rofA)*.

**Fig 7.**
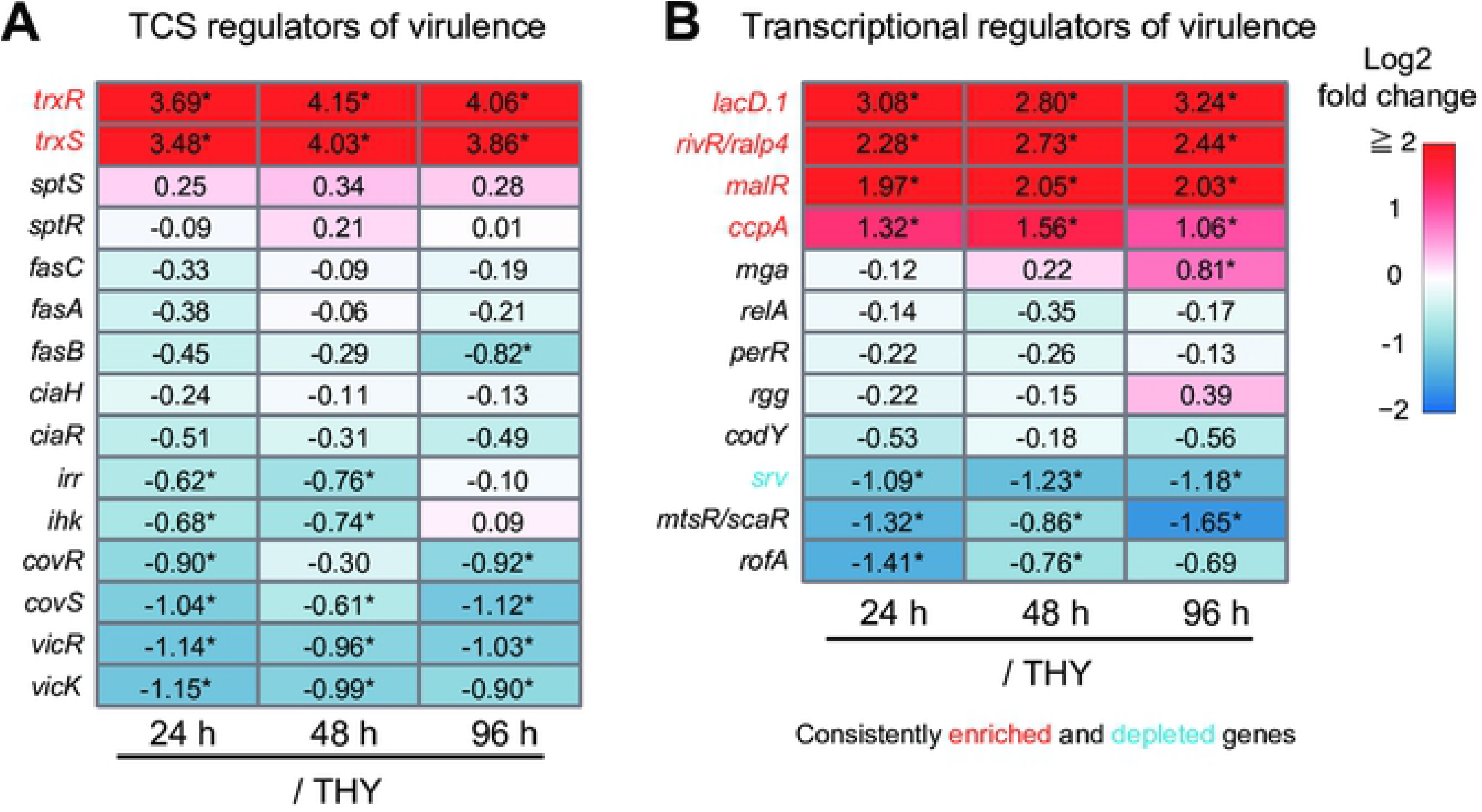
Expression levels of genes encoding virulence-related transcriptional regulators. (A) Two-component signal transduction systems (TCSs) and (B) transcriptional regulators of virulence. The log_2_ fold-changes of 24-h group (left), 48-h group (center), and 96-h group (right) with respect to THY group are indicated in color-scaled boxes. Numbers in the frame represent the measured log_2_ fold-change values. Color scale indicates enrichment (red) and depletion (blue) during infection. Asterisks indicate significant difference: *P < 0.1.

## Discussion

This is the first report of comprehensive gene-expression analyses of *S. pyogenes* in a mouse model of necrotizing fasciitis. For RNA-seq analysis of bacteria in host tissues, deep sequencing has been previously used to obtain a sufficient number of reads [37]. However, our protocol is simple and inexpensive and appears to effectively enable *in vivo* RNA-seq analysis of Gram-positive bacteria without deep sequencing. Here, we also analyzed the transcriptome profiles of *S. pyogenes* at three distinct time points during infection. Our results indicated that *S. pyogenes* drastically upregulates the expression of virulence-associated genes and shifts metabolic-pathway usage in the mouse model of necrotizing fasciitis consistently, and the results showed high-level expression in particular of *sagA*, *speB*, and *spd*. By contrast, *S. pyogenes* downregulated genes associated with oxidative-stress response and cell division in infected hind limbs relative to that in THY culture at the mid-logarithmic phase.

Our RNA-seq analysis revealed that *sagA, spi, speB* and *spd* were extremely upregulated in mouse necrotizing fasciitis as compared to in bacterial culture medium. Streptolysin S (SLS; encoded by *sagA-I*) and SpeB (encoded by *speB*) are widely recognized virulence factors of *S. pyogenes* [38]. SLS is involved in cellular injury, phagocytic resistance, and virulence in murine subcutaneous infection [39, 40], and SLS and SpeB promote *S. pyogenes* translocation via a paracellular route by degrading epithelial junctions [41, 42]. SpeB is a secreted cysteine protease degrading a wide variety of host proteins including complement components and cytokines, and functions in escape of *S. pyogenes* from host immune response [43–47]. Moreover, SpeB has been shown to contribute its virulence substantially in mouse models of necrotizing myositis [23, 48]. The *spi* and *speB* genes are co-transcribed [49]. The *spi* gene encodes a specific SpeB inhibitor, Spi, to protect bacterial cell from the activity of residual unsecreted SpeB. We found that DNases encoded by *sda1, spd3*, and *spd* were also upregulated markedly. Sda1 allows *S. pyogenes* to escape killing in neutrophil extracellular traps and contributes to virulence in murine subcutaneous infection [50, 51]. The expression of *spd*, which encodes streptodornase B or mitogenic factor 1, was ranked 4^th^ here, and a previous study has also reported its contribution to the virulence of *S. pyogenes* (serotype M89) [52]. Although *S. pyogenes* contains various virulence factors [12, 38, 53], these four genes showed outstanding up-regulation in our infection model. Our findings would help searching therapeutic targets for necrotizing facilities.

In this study, we also detected drastic upregulation of virulence genes encoding histidine triad protein (HtpA) [54] and laminin-binding protein (Lbp) [55]; our results could provide valuable insights regarding the utility of these molecules as therapeutic targets. Although we used a mouse intraperitoneal-infection model, we found that HtpA functions as an effective vaccine antigen against *S. pyogenes* [56]. Furthermore, analyses of sera from patients with uncomplicated *S. pyogenes* infection or rheumatic fever indicated the detectable humoral response against recombinant *S. pyogenes* Lbp [57].

The 483 genes that were consistently altered in this study overlap with the 150 low-glucose-induced genes of strain HSC5 (serotype 14) [17]. The overlapping genes include the upregulated genes encoding molecules involved in carbohydrate uptake and metabolism, arginine metabolism, V-Type ATP synthase, and lactate oxidase, whereas the overlapping downregulated genes contain molecules related to oxidative-stress response and cell division. In terms of the expression of genes encoding virulence factors, we observed the overlapping of upregulation of the genes for SLS, streptolysin O, and Spd and downregulation of GRAB gene. These findings suggest that *S. pyogenes* in the infected hindlimbs encounters a glucose-poor environment and relieves carbon catabolite repression [26].

Mutations in *covRS* of *S. pyogenes* serotype M1 (strains 5448) have been reported to enhance virulence during subcutaneous infection in mouse and might be responsible for loss of SpeB expression [51]. Graham *et al*. also reported that serotype M1 *S. pyogenes* (MGAS5005) showed reduced levels of the *speB* transcript during growth in human blood [21].

However, in this study, the gene encoding SpeB was drastically upregulated in the mouse model of necrotizing fasciitis (log_2_ fold-change > 9.38). The environment that *S. pyogenes* encounters in necrotizing fasciitis is considered to be distinct from that in blood and in subcutaneous tissue. Although blood pH is maintained in a narrow range around pH 7.4 in living organisms, inflammatory loci are typically associated with an acidic environment [58]. Moreover, our results suggested that *S. pyogenes* encounters glucose deprivation in necrotizing fasciitis. In *S. pyogenes, speB* expression at the early stationary phase can be substantially suppressed by glucose and buffered pH [59]. Generally, the stationary phase of bacterial growth is evidenced by glucose depletion and medium acidification. Thus, an environment similar to the bacterial stationary phase might have induced the strong expression of *speB*.

Graham *et al*. also characterized the MGAS5005 (serotype M1) transcript profile in a mouse soft-tissue infection model (subcutaneous infection) by using a wild-type strain and *ΔcovR* strain [20]; intriguingly, relative to the wild-type strain, *ΔcovR* strain exhibited drastic upregulation of *sagA* (18-fold), *speB* (2,053-fold), and *spd* (6-fold) in this model, and normalized expression levels of these 3 genes ranked 8^th^, 2^nd^, and 5^th^, respectively, in *ΔcovR* strain. In our study, *S. pyogenes* in the mouse model of necrotizing fasciitis also showed extremely high normalized expression levels of *sagA* (ranked 1^st^), *speB* (3^rd^), and *spd* (4^th^) among 1723 genes. One of the classic signs of acute inflammation is heat, and muscle temperature is considered to be higher than skin temperature [60]. *S. pyogenes* appears to encounter higher temperatures during myositis than during subcutaneous infection, which might lead to distinct transcriptome profiles of *S. pyogenes*.

Arginine and histidine are present in human muscle at high concentrations, ~1,000 and 500 μM, respectively [61]. Because a supply of amino acids is essential for protein and nucleic acid synthesis, the arginine and histidine metabolic pathways are likely to be enhanced, as was observed here, for pathogenicity to be exerted in necrotizing fasciitis. Moreover, for the uptake of essential amino acids, the operon encoding dipeptide transporter (DppABCDF) was consistently upregulated in the infected hindlimbs. Deletion of *S. pyogenes dppA* results in *speB* expression decreasing to one-eighth of its original level (serotype M49, strain CS101) [28]. Thus, *dppA* upregulation might contribute to the drastically increased expression of *speB*.

Recently, Zhu *et al*. identified the genes required for a cynomolgus macaque model of necrotizing fasciitis by using transposon-directed insertion-site sequencing [22]. The serotype M1 (MAGS2221) genes necessary for infection that were identified by Zhu *et al*. overlap with certain upregulated genes in our study, such as genes for carbohydrate metabolism (*glgP*, *malM*), arginine metabolism (*arcABCD*), and putative or known transporters (valine, *braB;* zinc, *adcBC;* SLS, *sagGHI)*. However, in transposon-directed insertion-site sequencing, insertion sites are detected after DNA-sequencing, implying that gene-expression levels are not considered. In RNA-seq analysis, relative expression levels among all genes can be evaluated. For the investigation of therapeutic targets, it is critical to select highly expressed molecules, which suggests the importance of our study for this purpose.

No transcriptome profiling of *S. pyogenes* in necrotizing fasciitis have been previously reported. This study revealed that *S. pyogenes* in the mouse model of necrotizing fasciitis exhibited substantially altered global transcription as compared to that under *in vitro* conditions. *S. pyogenes* might have attempted to acquire nutrients by destroying tissues by markedly upregulating the expression of toxins such as SLS, SpeB, and Spd. Furthermore, genes encoding molecules involved in carbohydrate and amino acid utilization as well as metal-transporter genes were upregulated in the infected mouse hindlimbs. We also believe that our protocol for isolating bacterial RNA from infected tissues at high concentrations will facilitate studies involving global gene-expression analyses of bacteria in the *in vivo* host environment. Future studies could explore new therapies based on bacterial kinetics *in vivo* by exploiting our data or our methods. The accumulation of *in vivo* gene-expression profiles will provide useful information necessary for establishing novel treatment strategies or identifying effective vaccine antigens.

## Materials and methods

### Ethic statement

All mouse experiments were conducted in accordance with animal protocols approved by the Animal Care and Use Committee of Osaka University Graduate School of Dentistry (28-002-0). Animals were cared for according to Guidelines for Proper Conduct of Animal Experiments (Science Council of Japan) and the policy laid down by the Animal Care and Use Committee of Osaka University Graduate School of Dentistry.

### Bacterial strains and culture conditions

*S. pyogenes* M1T1 strain 5448 (accession: CP008776) was isolated from a patient with toxic-shock syndrome and necrotizing fasciitis; the strain is genetically representative of a globally disseminated clone associated with invasive *S. pyogenes* infections [62]. *S. pyogenes* strain 5448 was cultured in Todd-Hewitt broth (BD Biosciences, San Jose, CA) supplemented with 0.2% yeast extract (BD Biosciences) (THY) at 37°C. For growth measurements, overnight cultures of *S. pyogenes* strain 5448 were back-diluted 1:50 into fresh THY and grown at 37°C; growth was monitored by measuring the optical density at 600 nm (OD_600_).

### Necrotizing fasciitis studies

We used 10-week-old male C57BL/6J mice (Charles River Japan Inc., Kanagawa, Japan) for the necrotizing fasciitis studies, as described previously [23]. After growing *S. pyogenes* cultures until the mid-exponential phase (OD_600_ = ~0.5), THY was replaced with PBS and the bacterial suspensions were stored in a refrigerator (−80°C). Viable cell counts of the suspensions were determined by plating diluted samples on THY blood agar. Mice were shaved and hair was removed through chemical depilation (Veet, Oxy Reckit Benckiser, Chartes, France), and then the mice were inoculated intramuscularly in both sides of hindlimbs with 2 × 10^7^ CFU suspended in 100 μL of PBS, prepared immediately before infection by diluting frozen stocks.

Mice were euthanized at 24, 48, or 96 h after infection by means of lethal intraperitoneal injection of sodium pentobarbital, and then the infected hindlimbs were collected. The left hindlimbs were immediately placed in RNAlater (Qiagen, Valencia, CA) and stored at −80°C until use in RNA isolation, whereas the right hindlimbs were fixed with formalin, embedded in paraffin and sectioned, and stained with hematoxylin and eosin, as described previously [63].

### RNA isolation

Thawed tissues were placed in lysing Matrix D microtubes containing 1.4-mm silica spheres (Qbiogene, Carlsbad, CA) with RLT lysis buffer (RNeasy Fibrous Tissue Mini Kit, Qiagen, Hilden, Germany) and homogenized at 6,500 rpm for 45 s by using a MagNA Lyser (Roche, Mannheim, Germany). The lysate was centrifuged, and the obtained pellet was resuspended in lysing Matrix B microtubes containing 0.1-mm silica spheres (Qbiogene) with the RLT lysis buffer and homogenized at 6,500 rpm for 60 s by using the MagNA Lyser. The final lysate was centrifuged, and bacterial RNA was isolated from the collected supernatant by using the RNeasy Fibrous Tissue Mini Kit according to the manufacturer’s guidelines and stored at −80°C (Fig 1B).

### RNA-seq and data analysis

RNA integrity was assessed using a 2100 Bioanalyzer (Agilent Technologies, Santa Clara, CA) (Fig 1C). For RNA-seq, 5 μg of bacterial RNA was treated for ribosomal RNA (rRNA) removal by using a Ribo-Zero rRNA Removal Kit (Mouse and Bacteria) (Illumina Inc., San Diego, CA). Directional RNA-seq libraries were created using TruSeq RNA Sample Prep Kit v2 (Illumina Inc.), according to the manufacturer’s recommendations. Libraries were sequenced using Illumina NovaSeq 6000 and HiSeq 2500 systems, with 100-bp paired-end reads being obtained (Macrogen, Daejeon, Korea). Data were generated in the standard Sanger FastQ format and raw reads were deposited into the DDBJ sequence read archive (DRA, accession number: DRA008246). Phred-type quality scores Q30 were used for quality trimming. RNA-seq reads were mapped against the *S. pyogenes* strain 5448 genome (accession CP008776) by using the commercially available CLC Genomics workbench (version 9.5.2, CLC Bio, Aarhus, Denmark). Differential expression analyses and global analysis of the RNA-seq expression data were performed using iDEP (http://ge-lab.org/idep/) [64], with the RPKM value of each sample being determined. Results were visualized using volcano plots (iDEP) and Venn diagrams (http://bioinformatics.psb.ugent.be/webtools/Venn/). EdgeR log-transformation was used for clustering and PCA (iDEP). The hierarchical clustering was illustrated by using the average-linkage method with correlation distance (iDEP). The data were also clustered by using k-means with 1,723 genes (k = 4) (iDEP). We classified the DEGs into functional categories based on the bacterial bioinformatics database and analysis resource PATRIC (www.patricbrc.org) [65], which is integrated with information from VFDB (http://www.mgc.ac.cn/VFs/) [66], Victors [67], subsystems technology toolkit (*RASTtk*) [68, 69], and KEGG map [70]. Genes were also classified into pathways based on BioCyc database [71]. The transcriptomic (RNA-seq) data are summarized in S1 Dataset.

## Acknowledgment

We acknowledge the NGS core facility of the Genome Information Research Center at the Research Institute for Microbial Diseases of Osaka University for the support in RNA sequencing and data analysis.

## Supporting information

**S1 Fig.**
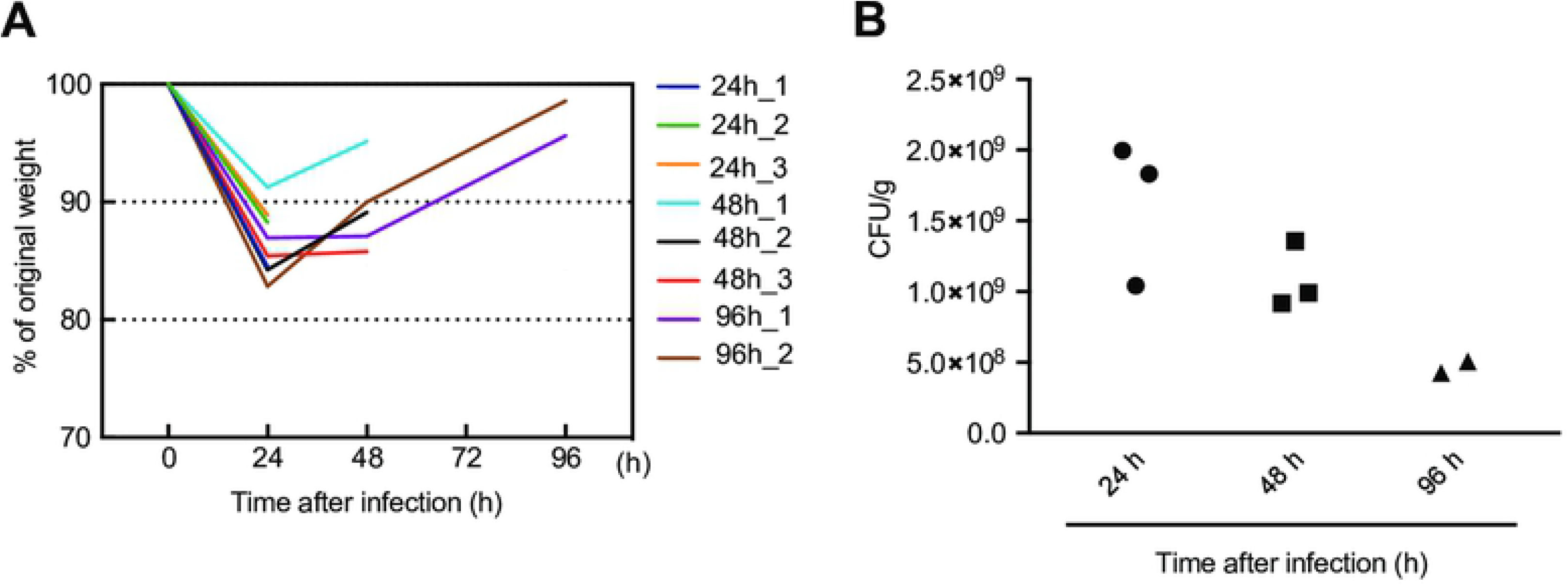
Mice recover from necrotizing fasciitis at 96 h after infection. Male C57BL/6J mice (10 weeks old) were intramuscularly inoculated in the hindlimbs with 2 × 10^7^ CFU of *S. pyogenes*. (A) Body-weight change until sample collection. Body weight at 0 h was regarded as 100%. (B) CFU of *S. pyogenes* in infected hindlimb samples.

**S2 Fig.**
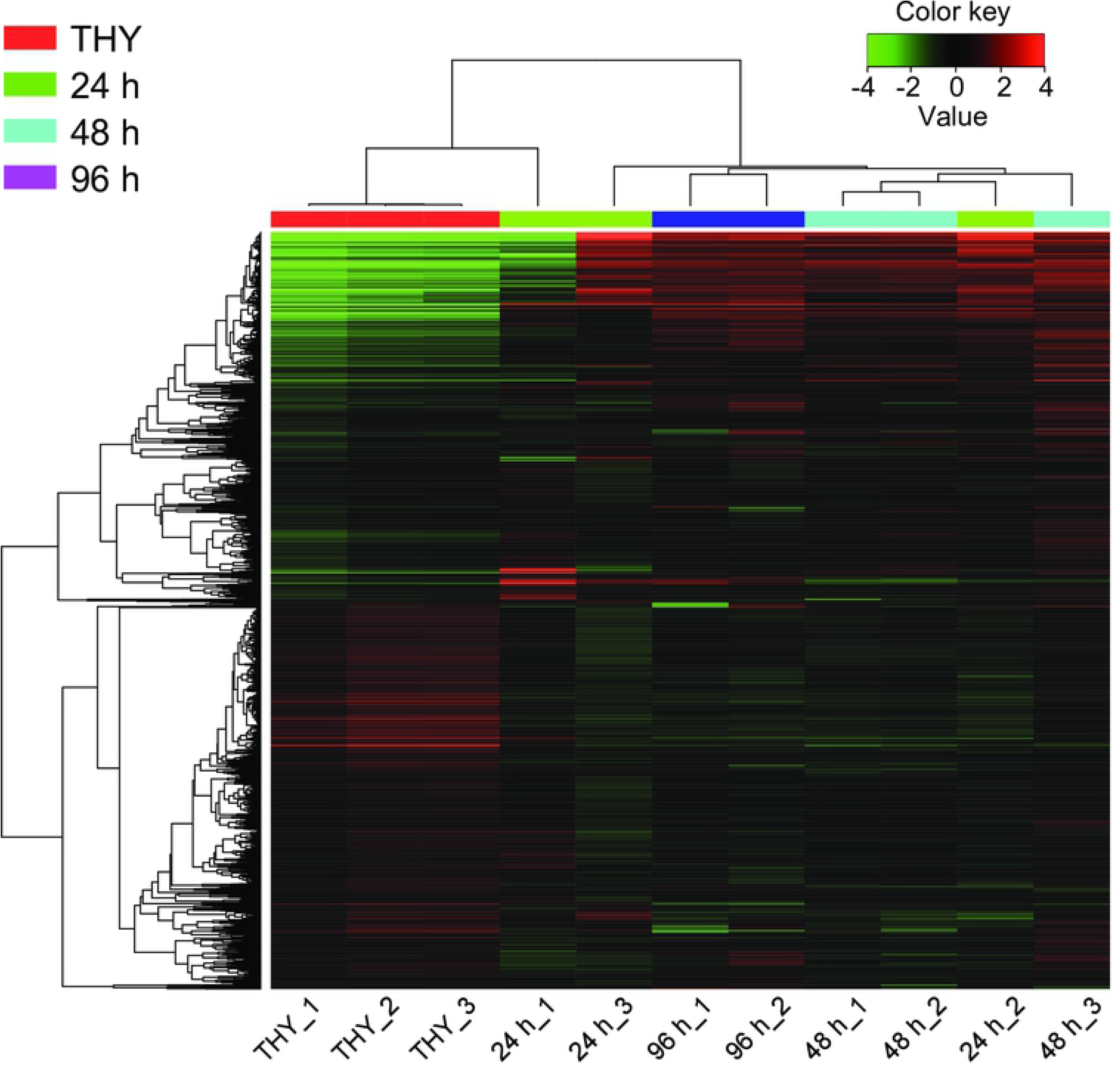
Heatmap of clustering of all genes (1,723 genes) expressed in all samples. Each column represents a sample, and each row represents a gene. Clustering was performed by using iDEP (http://ge-lab.org/idep/) with edgeR log-transformation of reads per kilobase million (RPKM) values. The hierarchical clustering was illustrated by using the average-linkage method with correlation distance. Color-coding is based on edgeR log-transformed RPKM values. The color key indicates the Z-scores, which display the relative values of all tiles within all samples: green, lowest expression; black, intermediate expression; red, highest expression.

**S1 Dataset. Global gene expression changes in a mouse model of necrotizing fasciitis.**

**S2 Dataset. Information of consistently altered mRNAs at three different distinct time-points during infection.**

## References

1. Stevens DL, Bryant AE. Severe Group A Streptococcal Infections. In: Ferretti JJ, Stevens DL, Fischetti VA, editors. Streptococcus pyogenes: Basic Biology to Clinical Manifestations. Oklahoma City (OK)2016.

2. Nelson GE, Pondo T, Toews KA, Farley MM, Lindegren ML, Lynfield R, et al. Epidemiology of Invasive Group A Streptococcal Infections in the United States, 2005-2012. Clin Infect Dis. 2016;63(4):478–86. doi: 10.1093/cid/ciw248. PMID: 27105747.

3. Misiakos EP, Bagias G, Patapis P, Sotiropoulos D, Kanavidis P, Machairas A. Current concepts in the management of necrotizing fasciitis. Front Surg. 2014;1:36. doi: 10.3389/fsurg.2014.00036. PMID: 25593960.

4. Carapetis JR, Steer AC, Mulholland EK, Weber M. The global burden of group A streptococcal diseases. Lancet Infect Dis. 2005;5(11):685–94. doi: 10.1016/S1473-3099(05)70267-X. PMID: 16253886.

5. Sanderson-Smith M, De Oliveira DM, Guglielmini J, McMillan DJ, Vu T, Holien JK, et al. A systematic and functional classification of *Streptococcus pyogenes* that serves as a new tool for molecular typing and vaccine development. J Infect Dis. 2014;210(8):1325–38. doi: 10.1093/infdis/jiu260. PMID: 24799598.

6. Steer AC, Law I, Matatolu L, Beall BW, Carapetis JR. Global *emm* type distribution of group A streptococci: systematic review and implications for vaccine development. Lancet Infect Dis. 2009;9(10):611–6. doi: 10.1016/S1473-3099(09)70178-1. PMID: 19778763.

7. Mora M, Bensi G, Capo S, Falugi F, Zingaretti C, Manetti AG, et al. Group A *Streptococcus* produce pilus-like structures containing protective antigens and Lancefield T antigens. Proc Natl Acad Sci U S A. 2005;102(43):15641–6. doi: 10.1073/pnas.0507808102. PMID: 16223875.

8. Falugi F, Zingaretti C, Pinto V, Mariani M, Amodeo L, Manetti AG, et al. Sequence variation in group A *Streptococcus* pili and association of pilus backbone types with lancefield T serotypes. J Infect Dis. 2008;198(12):1834–41. doi: 10.1086/593176. PMID: 18928376.

9. Ekelund K, Darenberg J, Norrby-Teglund A, Hoffmann S, Bang D, Skinhoj P, et al. Variations in *emm* type among group A streptococcal isolates causing invasive or noninvasive infections in a nationwide study. J Clin Microbiol. 2005;43(7):3101–9. doi: 10.1128/JCM.43.7.3101-3109.2005. PMID: 16000420.

10. Shea PR, Ewbank AL, Gonzalez-Lugo JH, Martagon-Rosado AJ, Martinez-Gutierrez JC, Rehman HA, et al. Group A *Streptococcus emm* gene types in pharyngeal isolates, Ontario, Canada, 2002-2010. Emerg Infect Dis. 2011;17(11):2010–7. doi: 10.3201/eid1711.110159. PMID: 22099088.

11. Ikebe T, Tominaga K, Shima T, Okuno R, Kubota H, Ogata K, et al. Increased prevalence of group A *streptococcus* isolates in streptococcal toxic shock syndrome cases in Japan from 2010 to 2012. Epidemiol Infect. 2015;143(4):864–72. doi: 10.1017/S0950268814001265. PMID: 25703404.

12. Walker MJ, Barnett TC, McArthur JD, Cole JN, Gillen CM, Henningham A, et al. Disease manifestations and pathogenic mechanisms of Group A *Streptococcus*. Clin Microbiol Rev. 2014;27(2):264–301. doi: 10.1128/CMR.00101-13. PMID: 24696436.

13. Aziz RK, Kotb M. Rise and persistence of global M1T1 clone of *Streptococcus pyogenes*. Emerg Infect Dis. 2008;14(10):1511–7. doi: 10.3201/eid1410.071660. PMID: 18826812.

14. Vega LA, Malke H, McIver KS. Virulence-Related Transcriptional Regulators of *Streptococcus pyogenes*. In: Ferretti JJ, Stevens DL, Fischetti VA, editors. Streptococcus pyogenes: Basic Biology to Clinical Manifestations. Oklahoma City (OK)2016.

15. Shelburne SA, Olsen RJ, Suber B, Sahasrabhojane P, Sumby P, Brennan RG, et al. A combination of independent transcriptional regulators shapes bacterial virulence gene expression during infection. PLoS Pathog. 2010;6(3):e1000817. doi: 10.1371/journal.ppat.1000817. PMID: 20333240.

16. Dalton TL, Collins JT, Barnett TC, Scott JR. RscA, a member of the MDR1 family of transporters, is repressed by CovR and required for growth of *Streptococcus pyogenes* under heat stress. J Bacteriol. 2006;188(1):77–85. doi: 10.1128/JB.188.1.77-85.2006. PMID: 16352823.

17. Kietzman CC, Caparon MG. Distinct time-resolved roles for two catabolite-sensing pathways during *Streptococcus pyogenes* infection. Infect Immun. 2011;79(2):812–21. doi: 10.1128/IAI.01026-10. PMID: 21098101.

18. Shelburne SA, Keith D, Horstmann N, Sumby P, Davenport MT, Graviss EA, et al. A direct link between carbohydrate utilization and virulence in the major human pathogen group A *Streptococcus*. Proc Natl Acad Sci U S A. 2008;105(5):1698–703. doi: 10.1073/pnas.0711767105. PMID: 18230719.

19. Kinkel TL, McIver KS. CcpA-mediated repression of streptolysin S expression and virulence in the group A *streptococcus*. Infect Immun. 2008;76(8):3451–63. doi: 10.1128/IAI.00343-08. PMID: 18490461.

20. Graham MR, Virtaneva K, Porcella SF, Gardner DJ, Long RD, Welty DM, et al. Analysis of the transcriptome of group A *Streptococcus* in mouse soft tissue infection. Am J Pathol. 2006;169(3):927–42. doi: 10.2353/ajpath.2006.060112. PMID: 16936267.

21. Graham MR, Virtaneva K, Porcella SF, Barry WT, Gowen BB, Johnson CR, et al. Group A *Streptococcus* transcriptome dynamics during growth in human blood reveals bacterial adaptive and survival strategies. Am J Pathol. 2005;166(2):455–65. doi: 10.1016/S0002-9440(10)62268-7. PMID: 15681829.

22. Zhu L, Olsen RJ, Beres SB, Eraso JM, Saavedra MO, Kubiak SL, et al. Gene fitness landscape of group A *streptococcus* during necrotizing myositis. J Clin Invest. 2019;129(2):887–901. doi: 10.1172/JCI124994. PMID: 30667377.

23. Olsen RJ, Sitkiewicz I, Ayeras AA, Gonulal VE, Cantu C, Beres SB, et al. Decreased necrotizing fasciitis capacity caused by a single nucleotide mutation that alters a multiple gene virulence axis. Proc Natl Acad Sci U S A. 2010;107(2):888–93. doi: 10.1073/pnas.0911811107. PMID: 20080771.

24. Olsen RJ, Musser JM. Molecular pathogenesis of necrotizing fasciitis. Annu Rev Pathol. 2010;5:1–31. doi: 10.1146/annurev-pathol-121808-102135. PMID: 19737105.

25. Keller N, Andreoni F, Reiber C, Luethi-Schaller H, Schuepbach RA, Moch H, et al. Human Streptococcal Necrotizing Fasciitis Histopathology Mirrored in a Murine Model. Am J Pathol. 2018;188(7):1517–23. doi: 10.1016/j.ajpath.2018.03.009. PMID: 29684366.

26. Deutscher J, Herro R, Bourand A, Mijakovic I, Poncet S. P-Ser-HPr--a link between carbon metabolism and the virulence of some pathogenic bacteria. Biochim Biophys Acta. 2005;1754(1-2):118–25. doi: 10.1016/j.bbapap.2005.07.029. PMID: 16182622.

27. Davies HC, Karush F, Rudd JH. Effect of Amino Acids on Steady-State Growth of a Group A Hemolytic *Streptococcus*. J Bacteriol. 1965;89:421–7. PMID: 14255710.

28. Podbielski A, Leonard BA. The group A streptococcal dipeptide permease (Dpp) is involved in the uptake of essential amino acids and affects the expression of cysteine protease. Mol Microbiol. 1998;28(6):1323–34. doi: 10.1046/j.1365-2958.1998.00898.x. PMID: 9680220.

29. Cusumano ZT, Watson ME, Jr., Caparon MG. *Streptococcus pyogenes* arginine and citrulline catabolism promotes infection and modulates innate immunity. Infect Immun. 2014;82(1):233–42. doi: 10.1128/IAI.00916-13. PMID: 24144727.

30. Brenot A, King KY, Caparon MG. The PerR regulon in peroxide resistance and virulence of *Streptococcus pyogenes*. Mol Microbiol. 2005;55(1):221–34. doi: 10.1111/j.1365-2958.2004.04370.x. PMID: 15612930.

31. King KY, Horenstein JA, Caparon MG. Aerotolerance and peroxide resistance in peroxidase and PerR mutants of *Streptococcus pyogenes*. J Bacteriol. 2000;182(19):5290–9. doi: 10.1128/JB.182.19.5290-5299.2000. PMID: 10986229.

32. Hanks TS, Liu M, McClure MJ, Fukumura M, Duffy A, Lei B. Differential regulation of iron- and manganese-specific MtsABC and heme-specific HtsABC transporters by the metalloregulator MtsR of group A *Streptococcus*. Infect Immun. 2006;74(9):5132–9. doi: 10.1128/IAI.00176-06. PMID: 16926405.

33. Weston BF, Brenot A, Caparon MG. The metal homeostasis protein, Lsp, of *Streptococcus pyogenes* is necessary for acquisition of zinc and virulence. Infect Immun. 2009;77(7):2840–8. doi: 10.1128/IAI.01299-08. PMID: 19398546.

34. Hood MI, Skaar EP. Nutritional immunity: transition metals at the pathogen-host interface. Nat Rev Microbiol. 2012;10(8):525–37. doi: 10.1038/nrmicro2836. PMID: 22796883.

35. Valdes KM, Sundar GS, Belew AT, Islam E, El-Sayed NM, Le Breton Y, et al. Glucose Levels Alter the Mga Virulence Regulon in the Group A *Streptococcus*. Sci Rep. 2018;8(1):4971. doi: 10.1038/s41598-018-23366-7. PMID: 29563558.

36. Churchward G. The two faces of Janus: virulence gene regulation by CovR/S in group A streptococci. Mol Microbiol. 2007;64(1):34–41. doi: 10.1111/j.1365-2958.2007.05649.x. PMID: 17376070.

37. Nuss AM, Beckstette M, Pimenova M, Schmühl C, Opitz W, Pisano F, et al. Tissue dual RNA-seq allows fast discovery of infection-specific functions and riboregulators shaping host-pathogen transcriptomes. Proc Natl Acad Sci U S A. 2017;114(5):E791–E800. doi: 10.1073/pnas.1613405114. PMID: 28096329.

38. Hamada S, Kawabata S, Nakagawa I. Molecular and genomic characterization of pathogenic traits of group A *Streptococcus pyogenes*. Proc Jpn Acad Ser B Phys Biol Sci. 2015;91(10):539–59. doi: 10.2183/pjab.91.539. PMID: 26666305.

39. Humar D, Datta V, Bast DJ, Beall B, De Azavedo JC, Nizet V. Streptolysin S and necrotising infections produced by group G *streptococcus*. Lancet. 2002;359(9301):124–9. doi: 10.1016/S0140-6736(02)07371-3. PMID: 11809255.

40. Datta V, Myskowski SM, Kwinn LA, Chiem DN, Varki N, Kansal RG, et al. Mutational analysis of the group A streptococcal operon encoding streptolysin S and its virulence role in invasive infection. Mol Microbiol. 2005;56(3):681–95. doi: 10.1111/j.1365-2958.2005.04583.x. PMID: 15819624.

41. Sumitomo T, Nakata M, Higashino M, Jin Y, Terao Y, Fujinaga Y, et al. Streptolysin S contributes to group A streptococcal translocation across an epithelial barrier. J Biol Chem. 2011;286(4):2750–61. doi: 10.1074/jbc.M110.171504. PMID: 21084306.

42. Sumitomo T, Mori Y, Nakamura Y, Honda-Ogawa M, Nakagawa S, Yamaguchi M, et al. Streptococcal Cysteine Protease-Mediated Cleavage of Desmogleins Is Involved in the Pathogenesis of Cutaneous Infection. Front Cell Infect Microbiol. 2018;8:10. doi: 10.3389/fcimb.2018.00010. PMID: 29416987.

43. von Pawel-Rammingen U, Björck L. IdeS and SpeB: immunoglobulin-degrading cysteine proteinases of *Streptococcus pyogenes*. Curr Opin Microbiol. 2003;6(1):50–5. doi: 10.1016/S1369-5274(03)00003-1. PMID: 12615219.

44. Honda-Ogawa M, Ogawa T, Terao Y, Sumitomo T, Nakata M, Ikebe K, et al. Cysteine proteinase from *Streptococcus pyogenes* enables evasion of innate immunity via degradation of complement factors. J Biol Chem. 2013;288(22):15854–64. doi: 10.1074/jbc.M113.469106. PMID: 23589297.

45. Terao Y, Mori Y, Yamaguchi M, Shimizu Y, Ooe K, Hamada S, et al. Group A streptococcal cysteine protease degrades C3 (C3b) and contributes to evasion of innate immunity. J Biol Chem. 2008;283(10):6253–60. doi: 10.1074/jbc.M704821200. PMID: 18160402.

46. Kapur V, Majesky MW, Li LL, Black RA, Musser JM. Cleavage of interleukin 1β (IL-1β) precursor to produce active IL-1β by a conserved extracellular cysteine protease from *Streptococcus pyogenes*. Proc Natl Acad Sci U S A. 1993;90(16):7676–80. doi: 10.1073/pnas.90.16.7676. PMID: 7689226.

47. Chiang-Ni C, Wu JJ. Effects of streptococcal pyrogenic exotoxin B on pathogenesis of *Streptococcus pyogenes*. J Formos Med Assoc. 2008;107(9):677–85. doi: 10.1016/S0929-6646(08)60112-6. PMID: 18796357.

48. Do H, Makthal N, VanderWal AR, Rettel M, Savitski MM, Peschek N, et al. Leaderless secreted peptide signaling molecule alters global gene expression and increases virulence of a human bacterial pathogen. Proc Natl Acad Sci U S A. 2017;114(40):E8498–E507. doi: 10.1073/pnas.1705972114. PMID: 28923955.

49. Kagawa TF, O’Toole P W, Cooney JC. SpeB-Spi: a novel protease-inhibitor pair from *Streptococcus pyogenes*. Mol Microbiol. 2005;57(3):650–66. doi: 10.1111/j.1365-2958.2005.04708.x. PMID: 16045611.

50. Buchanan JT, Simpson AJ, Aziz RK, Liu GY, Kristian SA, Kotb M, et al. DNase expression allows the pathogen group A *Streptococcus* to escape killing in neutrophil extracellular traps. Curr Biol. 2006;16(4):396–400. doi: 10.1016/j.cub.2005.12.039. PMID: 16488874.

51. Walker MJ, Hollands A, Sanderson-Smith ML, Cole JN, Kirk JK, Henningham A, et al. DNase Sda1 provides selection pressure for a switch to invasive group A streptococcal infection. Nat Med. 2007;13(8):981–5. doi: 10.1038/nm1612. PMID: 17632528.

52. Sriskandan S, Unnikrishnan M, Krausz T, Cohen J. Mitogenic factor (MF) is the major DNase of serotype M89 *Streptococcus pyogenes*. Microbiology. 2000;146 (Pt 11):2785–92. doi: 10.1099/00221287-146-11-2785. PMID: 11065357.

53. Yamaguchi M, Terao Y, Kawabata S. Pleiotropic virulence factor – *Streptococcus pyogenes* fibronectin-binding proteins. Cell Microbiol. 2013;15(4):503–11. doi: 10.1111/cmi.12083. PMID: 23190012.

54. Wen YT, Wang JS, Tsai SH, Chuan CN, Wu JJ, Liao PC. Label-free proteomic analysis of environmental acidification-influenced *Streptococcus pyogenes* secretome reveals a novel acid-induced protein histidine triad protein A (HtpA) involved in necrotizing fasciitis. J Proteomics. 2014;109:90–103. doi: 10.1016/j.jprot.2014.06.026. PMID: 24998435.

55. Terao Y, Kawabata S, Kunitomo E, Nakagawa I, Hamada S. Novel laminin-binding protein of *Streptococcus pyogenes*, Lbp, is involved in adhesion to epithelial cells. Infect Immun. 2002;70(2):993–7. doi: 10.1128/IAI.70.2.993-997.2002. PMID: 11796638.

56. Kunitomo E, Terao Y, Okamoto S, Rikimaru T, Hamada S, Kawabata S. Molecular and biological characterization of histidine triad protein in group A streptococci. Microbes Infect. 2008;10(4):414–23. doi: 10.1016/j.micinf.2008.01.003. PMID: 18403236.

57. Wahid RM, Yoshinaga M, Nishi J, Maeno N, Sarantuya J, Ohkawa T, et al. Immune response to a laminin-binding protein (Lmb) in group A streptococcal infection. Pediatr Int. 2005;47(2):196–202. doi: 10.1111/j.1442-200x.2005.02038.x. PMID: 15771700.

58. Lardner A. The effects of extracellular pH on immune function. J Leukoc Biol. 2001;69(4):522–30. doi: 10.1189/jlb.69.4.522. PMID: 11310837.

59. Loughman JA, Caparon M. Regulation of SpeB in *Streptococcus pyogenes* by pH and NaCl: a model for *in vivo* gene expression. J Bacteriol. 2006;188(2):399–408. doi: 10.1128/JB.188.2.399-408.2006. PMID: 16385029.

60. Costello JT, Culligan K, Selfe J, Donnelly AE. Muscle, skin and core temperature after – 110°C cold air and 8°C water treatment. PLoS One. 2012;7(11):e48190. doi: 10.1371/journal.pone.0048190. PMID: 23139763.

61. Canepa A, Filho JC, Gutierrez A, Carrea A, Forsberg AM, Nilsson E, et al. Free amino acids in plasma, red blood cells, polymorphonuclear leukocytes, and muscle in normal and uraemic children. Nephrol Dial Transplant. 2002;17(3):413–21. doi: 10.1093/ndt/17.3.413. PMID: 11865086.

62. Kansal RG, McGeer A, Low DE, Norrby-Teglund A, Kotb M. Inverse relation between disease severity and expression of the streptococcal cysteine protease, SpeB, among clonal M1T1 isolates recovered from invasive group A streptococcal infection cases. Infect Immun. 2000;68(11):6362–9. doi: 10.1128/IAI.68.11.6362-6369.2000. PMID: 11035746.

63. Hirose Y, Yamamoto T, Nakashima M, Funahashi Y, Matsukawa Y, Yamaguchi M, et al. Injection of Dental Pulp Stem Cells Promotes Healing of Damaged Bladder Tissue in a Rat Model of Chemically Induced Cystitis. Cell Transplant. 2016;25(3):425–36. doi: 10.3727/096368915X689523. PMID: 26395427.

64. Ge SX, Son EW, Yao R. iDEP: an integrated web application for differential expression and pathway analysis of RNA-Seq data. BMC Bioinformatics. 2018;19(1):534. doi: 10.1186/s12859-018-2486-6. PMID: 30567491.

65. Wattam AR, Davis JJ, Assaf R, Boisvert S, Brettin T, Bun C, et al. Improvements to PATRIC, the all-bacterial Bioinformatics Database and Analysis Resource Center. Nucleic Acids Res. 2017;45(D1):D535–D42. doi: 10.1093/nar/gkw1017. PMID: 27899627.

66. Liu B, Zheng D, Jin Q, Chen L, Yang J. VFDB 2019: a comparative pathogenomic platform with an interactive web interface. Nucleic Acids Res. 2019;47(D1):D687–D92. doi: 10.1093/nar/gky1080. PMID: 30395255.

67. Sayers S, Li L, Ong E, Deng S, Fu G, Lin Y, et al. Victors: a web-based knowledge base of virulence factors in human and animal pathogens. Nucleic Acids Res. 2019;47(D1):D693–D700. doi: 10.1093/nar/gky999. PMID: 30365026.

68. Aziz RK, Bartels D, Best AA, DeJongh M, Disz T, Edwards RA, et al. The RAST Server: rapid annotations using subsystems technology. BMC Genomics. 2008;9:75. doi: 10.1186/1471-2164-9-75. PMID: 18261238.

69. Brettin T, Davis JJ, Disz T, Edwards RA, Gerdes S, Olsen GJ, et al. RASTtk: a modular and extensible implementation of the RAST algorithm for building custom annotation pipelines and annotating batches of genomes. Sci Rep. 2015;5:8365. doi: 10.1038/srep08365. PMID: 25666585.

70. Kanehisa M, Sato Y, Kawashima M, Furumichi M, Tanabe M. KEGG as a reference resource for gene and protein annotation. Nucleic Acids Res. 2016;44(D1):D457–62. doi: 10.1093/nar/gkv1070. PMID: 26476454.

71. Caspi R, Altman T, Billington R, Dreher K, Foerster H, Fulcher CA, et al. The MetaCyc database of metabolic pathways and enzymes and the BioCyc collection of Pathway/Genome Databases. Nucleic Acids Res. 2014;42(Database issue):D459–71. doi: 10.1093/nar/gkt1103. PMID: 24225315.

